# Vec2SPARQL: integrating SPARQL queries and knowledge graph embeddings

**DOI:** 10.1101/463778

**Authors:** Maxat Kulmanov, Senay Kafkas, Andreas Karwath, Alexander Malic, Georgios V Gkoutos, Michel Dumontier, Robert Hoehndorf

**Affiliations:** Computer, Electrical and Mathematical Science and Engineering Division, Computational Bioscience Research Center, King Abdullah University of Science and Technology, Thuwal 23955, Saudi Arabia; Centre for Computational Biology, University of Birmingham, Birmingham, UK; Institute of Data Science, Maastricht University, Maastricht, The Netherlands

**Keywords:** SPARQL, vector space, knowledge graph

## Abstract

Recent developments in machine learning have lead to a rise of large number of methods for extracting features from structured data. The features are represented as a vectors and may encode for some semantic aspects of data. They can be used in a machine learning models for different tasks or to compute similarities between the entities of the data. SPARQL is a query language for structured data originally developed for querying Resource Description Framework (RDF) data. It has been in use for over a decade as a standardized NoSQL query language. Many different tools have been developed to enable data sharing with SPARQL. For example, SPARQL endpoints make your data interoperable and available to the world. SPARQL queries can be executed across multiple endpoints. We have developed a Vec2SPARQL, which is a general framework for integrating structured data and their vector space representations. Vec2SPARQL allows jointly querying vector functions such as computing similarities (cosine, correlations) or classifications with machine learning models within a single SPARQL query. We demonstrate applications of our approach for biomedical and clinical use cases. Our source code is freely available at https://github.com/bio-ontology-research-group/vec2sparql and we make a Vec2SPARQL endpoint available at http://sparql.bio2vec.net/.

## 1 Introduction

SPARQL is a standardized NoSQL query language originally developed for data represented in the Resource Description Framework (RDF) [36] format. It supports basic inference on graphs and can be used to query multiple endpoint simultaneously. The flexibility of SPARQL has led the development of a large number of adapters and wrappers that enable querying data formats and storage technologies beyond RDF, including SQL databases [29], text files and large tables [10], or any other kind of *structured* data.

The flexibility and wide applicability of SPARQL makes it well suited for managing large heterogeneous data infrastructures such as found in many clinics and hospitals [35], or complex research data infrastructures such as UniProt [42] and other genomic databases [18].

SPARQL is well-suited for querying *structured* data, including metadata. However, it is not well suited to query the “content” of unprocessed and unstructured data such as images, videos, or large text corpora. For example, querying all chest x-ray images in a database that show a cardiomegaly in male patients is possible using SPARQL based on the metadata attached to the images but cannot be done based on the content of the images alone. Methods that could determine whether an x-ray image shows such a phenotype will commonly rely on extracting features from an image and using a machine learning algorithm.

Recently, several machine learning methods have been developed that extract features from unstructured data. These features are represented within a vector space and may encode some aspects of semantics. Deep learning techniques have been applied to images [34], audio [14], video [19], but also to domain-specific data types such as protein sequences [21] or DNA [45]. Deep Learning models can also be applied to structured datasets such as RDF itself [2, 32], or to formal knowledge bases and ontologies [38]. To collect domain-specific deep learning models, domain specific model repositories have been developed in which these models are shared and made reusable and interoperable [6].

The vector space feature representations generated from deep learning models can also be queried using functions that perform vector operations and may have some defined semantics. For example, similarity measures such as cosine similarity, correlation coefficients between vectors, or other similarity measures are sometimes used to determine relatedness between entities from which features were extracted [24, 25], and certain vector transformation may be used for analogical reasoning and inference [7, 8].

While the vector space representations resulting from machine learning systems enable a type of query on a dataset, the kinds of queries that can be asked about vector space representations are largely disconnected from SPARQL. SPARQL is applicable to querying of structured data and meta-data, while vector operations can identify semantic relatedness based on features extracted from an item. It can be useful to combine queries involving semantic relatedness (with respect to a particular feature extraction model) and structured information represented as meta-data. For example, once feature vectors are extracted from images, meta-data that is associated with the images (such as geo-locations, image types, author, or similar) could be queried using SPARQL and *combined* with the semantic queries over the feature vectors extracted from the images themselves. Such a combination would, for example, allow to identify the images authored by person *a* that are most similar to an image of author *b*; it can enable similarity- or analogy-based search and retrieval in precisely delineated subsets; or, when feature learning is applied to structured datasets, can combine similarity search and link prediction based on knowledge graph embeddings with structured queries based on SPARQL.

Here, we present a general framework to integrate vector space representations of data together with their metadata, and query both within a joint framework. We implement a prototype of this framework using (a) a repository of feature vector representations associated with entities in a knowledge graph, (b) a mapping between the entities in the repository and the knowledge graph, (c) a method to retrieve entities from the repository based on vector space operations, among a specified set of entities, and (d) a set of function extensions for SPARQL that make this search accessible from within a SPARQL query and semantically integrate these operations with the SPARQL syntax and semantics. We make our prototype implementation as well as a demo freely available on Github ^1^. Furthermore, we demonstrate using biomedical, clinical, and bioinformatics use cases how our approach can enable new kinds of queries and applications that combine symbolic processing and retrieval of information through sub-symbolic semantic queries within vector spaces.

## 2 Vector space projections and operations

A large number of machine learning models have been developed that can take data of various types as input and project them onto vectors that capture or represent some aspects of the semantics within a vector space. A large number of models are available for text [24], knowledge graphs [2, 7, 32], images (x-ray, dermascope, etc.) [30, 12], and audio [14].

In the case of RDF and OWL data, the vector space projection can be done either trivially by representing classes as binary vectors or through machine learning. As an example for the first case, a type of genes could be represented as a binary vector based on its associated GO classes to be further used in a similarity measurement such as a wide range of semantic similarity measures [13, 28] or vector similarity measures, to determine functionally similar genes. For more complex data, however, machine learning algorithm can now be used to extract relevant features and generate vector representations of nodes and relations in knowledge graphs. As these vector aim to encode information about nodes that represent the local structure in which a node is embedded, these vectors are called knowledge graph embeddings.

For example, these embeddings can be generated based on a random walk approach in combination with word2vec [26, 2] to generate embeddings for diseases, genes, or drugs [1], and which have shown utility in predicting gene-disease associations. Other approaches rely on constrained optimization where certain invariances that exist in a knowledge graph are preserved with respect to certain vector space operations. For example, Translational Embeddings (TransE) [7] which represent knowledge graph relations as translations in a vector space. A number of translational embedding methods have been proposed which are based on TransE [44, 16, 17]

Once the biomedical entities are represented as vectors, different methods can be applied to measure the similarity between the entities. The most widely used similarity measures for vectors generated as word or knowledge graph embeddings are cosine similarity, and, for many types of feature vectors, also correlation coefficients (e.g., Pearson and Spearman correlation) which are often used to compare features and identify similarity between entities.

Cosine similarity measures the orientation of two *n*-dimensional vectors. It is calculated by the dot product of two numeric vectors, and it is normalized by the dot product of the vectors’ magnitudes. Output values close to 1 indicate high similarity. Cosine similarity between two vectors *x* and *y* is formally defined in equation 1 as follows:

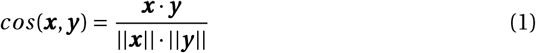

The Pearson correlation coefficient is used to measure linear relationships between two variables. An output value of 1 represents a perfect positive correlation, *−*1 indicates a perfect negative relationship, and 0 indicates the absence of a relationship between variables. Peason’s correlation coefficient of *X* and *Y* is formulated as:

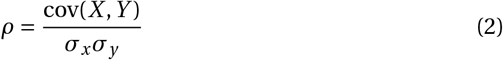

However, in general, any function *f*: ℝ*n ×*: ℝ*n •→*: ℝ that takes two vectors as input and outputs a real number can be used to determine similarity between vectors and potentially measure a form of “semantic” similarity between the entities represented by the vectors. Specifically, artificial neural networks (ANNs) can implement (or approximate) any function [15] and can be trained to output specific types of similarity, for example using a sigmoid classification function as output.

Computing similarity is not the only operation that can be performed on vector spaces. Some vector space models are built to preserve certain semantic invariances under other operations, such as addition and subtraction of vectors. For example, analogical reasoning can be performed using addition and subtraction based on word embeddings generated by Word2Vec [24], and in translational embedding models it is possible to add relation vectors to vectors representing nodes in the graph to perform multi-relational link prediction [7, 44, 16, 17].

## 3 Vec2SPARQL: jointly querying structured data and vector space representations

Vec2SPARQL bridges vector space embeddings of entities and the structured data about the entities that are accessible through SPARQL in a single framework. Vec2SPARQL assumes on the vector space side the existence of a repository (or a set) of vectors and the ability to perform certain vector space operations (such as computing similarity between two vectors). Each vector representation must be identified through an Internationalized Resource Identifier (IRI). The vectors, their IRIs, and the vector space operations should be made available through an API. On the SPARQL side, Vec2SPARQL assumes the existence of a SPARQL end-point linked to either structured data (e.g. knowledge graph) or semi-structured data (e.g., text documents). In SPARQL, entities and relations are identified through an IRI. Vec2SPARQL assumes that there is an overlap between the IRIs accessible through the SPARQL endpoint and the IRIs that are used to identify the vectors. The underlying idea is that the structured and semi-structured information about an entity can be queried through SPARQL, and the unstructured information about the entity can be queried through the vector space representations. For example, we can identify metadata about chest x-ray images in SPARQL, including an IRI to identify the image, the date the image was taken, the anatomical location, the diagnosis, and other meta-data relating to a patient; then, using a feature extractor for x-ray images, we can generate a vector representation and identify the vector with the same IRI as the image, and use vector space similarity (e.g., Pearson correlation coefficients) to identify similar or related images. Vec2SPARQL enables a bi-directional information flow between both types of information and combinations of these queries.

Specifically, in Vec2SPARQL, we extend the SPARQL query syntax by adding two custom functions with a particular semantics. The first function is *similarity*(?*x*,?*y*) where *x* and *y* are entities that are identified by an IRI in SPARQL. The function computes similarity of the corresponding embedding vectors. *similarity* is an expression function and can be used in FILTER, BIND, and SELECT statements.

The second function in Vec2SPARQL is called *mostSimilar*(?*x*, *n*) where *x* is an IRI of an entity accessible through SPARQL and for which a vector representation exists, and *n* is an integer. *most Similar* is a property function that allows to create new matches within a query using a similarity function define on the vector representations of entities. As a result of this function we will get *n* entities which are the most similar to *x* and can further be processed with query operators.

Vec2SPARQL is implemented using ARQ, which is a query engine for Apache Jena [9]. It currently implements only a single similarity function (cosine similarity) but can easily be extended to other functions. When multiple different similarity functions are to be used, we will extend the Vec2SPARQL functions with an additional argument that specifies which function to use.

Vec2SPARQL can easily be run using a Docker [23]. Docker is an open-source container software which allows to distribute, deploy and run a software tools in a virtual environment. We have configured a docker image which installs all required dependencies and builds the Vec2SPARQL executable in order to start the SPARQL endpoint.

## 4 Remote querying of vector similarity

To build a more flexible infrastructure we do not maintain the repository of vector representations or compute the similarity functions directly in Vec2SPARQL but rather use an API that consists of a repository of such vectors and a set of functions (mainly search by similarity) to execute on them. For this purpose, we rely on the prototypical Bio2Vec platform. Bio2Vec is a platform for representing, sharing, integrating, and querying vector space embeddings. Its current content covers embeddings from text, knowledge graphs, and biological interaction networks, and has vector representations of several types of biological and biomedical entities, including gene functions (from Gene Ontology [5]), genes, drugs, and diseases. Bio2Vec has an API through which the vector space can be searched by similarity, currently using only the cosine similarity measure.

Vec2SPARQL utilizes Bio2Vec and its similarity functions within SPARQL queries. We implemented a first version of Vec2SPARQL to work with Bio2Vec API. However, Vec2SPARQL can be integrated with other external APIs which provide similar kinds of vector space operations.

Figure 1 illustrates the general picture of the Vec2SPARQL approach. Vec2SPARQL provides a single SPARQL endpoint in which queries can be performed over the Bio2Vec API together with data currently stored in the Vec2SPARQL endpoint. We use the Bio2Vec REST API, which is based on an ElasticSearch index with a vector scoring plugin, to store and search for embedding vectors. When a Vec2SPARQL custom function is used in a query, Vec2SPARQL retrieves embeddings or similarity values from Bio2Vec. On the other side, the structured data is represented as RDF and is queried using Apache Jena’s ARQ query engine.

**Fig. 1.**
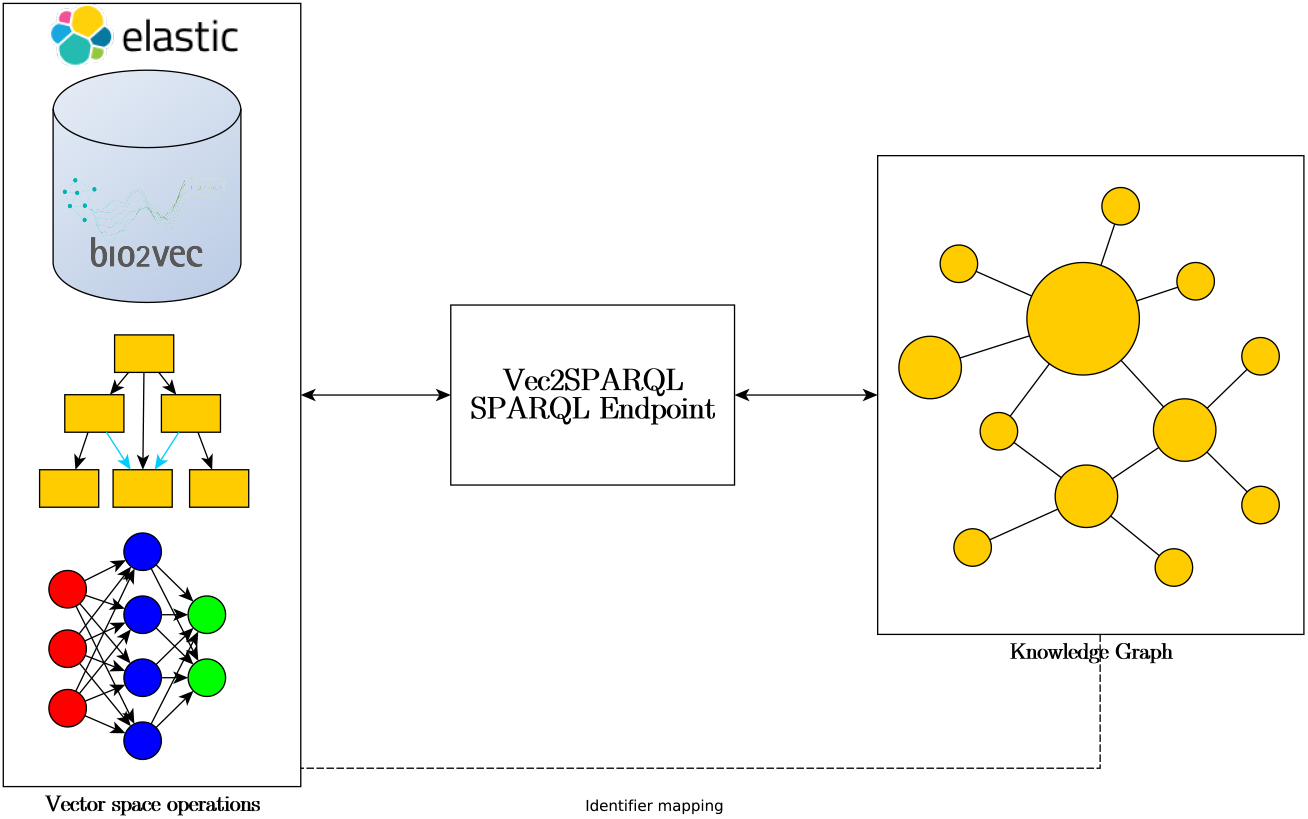
Overview over the Vec2SPARQL approach. Vec2SPARQL provides a SPARQL endpoint which can query a structured dataset using SPARQL and simultaneously perform vector space operations for the entities. Vec2SPARQL requires that there is a mapping between identifiers used to identify vectors and entities retrieved through SPARQL.

## 5 Use case: phenotype-driven disease gene prioritization

One widely used application of similarity-based search in biomedical applications is the comparison of phenotypes using a similarity measure. Recently, several vector space models have been developed that show competitive performance when using phenotype similarity within and across species to predict gene–disease associations [1, 39]. In these models, data is prepared in a custom manner and vector similarity (or, in some cases, a neural network) is used to exhaustively compute similarity between genes and diseases. We can use Vec2SPARQL to perform queries of a knowledge graph of mouse genes, diseases and phenotypes and incorporate Vec2SPARQL similarity functions.

We created a knowledge graph of mouse genes and their phenotype annotations obtained from the Mouse Genome Informatics (MGI) [40] database (accessed on 06.08.2018) and human diseases in the Online Mendelian Inheritance in Men (OMIM) [3] database, and their phenotype associations from the Human Phenotype Ontology (HPO) database (accessed on 27.07.2018). Our aim in this use case is to find mouse gene associations with human diseases by prioritizing them using their phenotypic similarity, and simultaneously restrict the similarity comparisons to genes and diseases with specific properties (such as being associated with a particular phenotype). The phenotypic similarity can be computed in different ways. One way is to use ontology based semantic similarity measure such as simGIC [27] or Resnik similarity measure [31]. However, there are hundreds of similarity measures and it is difficult to choose one. Also, semantic similarity measures can be biased towards well studied genes when the variance of annotation size is very high [20].

Another way of computing similarity is to use representation learning methods which can provide an embedding vector for the entities of a dataset. The embeddings can further be used as features for machine learning methods or compute similarities such as cosine or correlation coefficients depending on type of the embeddings.

Here, we employ a knowledge graph representation learning method by [2]. To include semantics of phenotype annotations both from Mammalian Phenotype Ontology (MP) and Human Phenotype Ontology (HPO) we add an integrated phenotype ontology PhenomeNET [33] to our knowledge graph. First, the method generates a corpus by randomly walking the knowledge graph including edge information. The corpus captures a neighborhood of each entity in the knowledge graph and can further be used in representation learning methods. Then, we use Word2Vec [24] and extract embeddings vectors for each entity in our knowledge graph. The Word2Vec embeddings are optimized for a cosine similarity. In other words, we can get similar entities of our knowledge graph by computing the cosine similarity of their embedding vectors.

Our hypothesis is that associated genes and diseases should have similar phenotype associations, therefore they should have more similar embeddings than not associated genes and diseases. Using Vec2SPARQL, we can answer this question with a simple SPARQL query. For example, the following query extracts disease associations for a mouse gene *Pax6* (MGI:97490):

~~~
PREFIX b2v: <http://bio2vec.net/graph_embeddings/function#> PREFIX MGI: <http://www.informatics.jax.org/gene/MGI_>
PREFIX obo: <http://purl.obolibrary.org/>
PREFIX rdf: <http://www.w3.org/1999/02/22-rdf-syntax-ns#>
PREFIX rdfs: <http://www.w3.org/2000/01/rdf-schema#>
SELECT ?sim ?dis (b2v:similarity(?sim, MGI:97490) as ?val)
{
?sim b2v:mostSimilar(MGI:97490 10000) .
?sim a obo:disease .
?sim rdfs:label ?dis
}
~~~

This query will return all diseases in OMIM among first 10,000 similar entities in the knowledge graph ordered by the cosine similarity value.

## 6 Use case: image repositories in health care environments

During decision making in the medical and clinical domains, physicians and consultants are often required to not only base their diagnosis on their medical expert knowledge, but furthermore to relate this diagnosis to past patients. To extract and model medical and clinical expert knowledge, a number of approaches based on machine learning and artificial intelligence have been proposed in the last decades. In recent years, this trend has been even more accelerated with the rise of deep convolutional networks. Some examples include the segmentation of organs and tissues in medical images or the prediction of diagnoses [22] and referral urgency using 3D Optical coherence tomography (OCT) images [11]. Other approaches aim to identify similar case histories to enhance the cased-based retrieval of similar patients based on the available medical information, often captured using images [4, 37].

Here, we employ an approach inspired by these last approaches in order to demonstrate the general applicability of Vec2SPARQL within the medical and clinical domains. We employed a publicly available dataset of chest x-ray images [43] which is also available from Kaggle for use in machine learning and analytics challenges. It consist of more than 112,000 chest x-ray images including basic annotations such as gender, age, and diagnosis of the patients. We downloaded the images and their annotations (accessed on 12.08.2018)) and scaled the original images to a resolution of 256 *×* 256 pixels. We then used the BVLC GoogLeNet [41] from the Caffe Model Zoo to extract feature vectors from these images. Specifically, we presented each re-sized image to the network and extracted the last layer before the final softmax layer (pool5/7 × 7_s1) as vector for use in Vec2SPARQL. Within our system, we have employed a set of approximately randomly chosen 15,881 images from a variety of medical diagnoses, patient ages, and gender.

Our hypothesis is that similar x-ray images should have similar clinical diagnoses. Using Vec2SPARQL, we can evaluate this with a SPARQL query. The following query for example extracts similar chest x-ray images to a patient with patient ID 9890, represented by the image 00009890_001.png (patient ID 9890, diagnosis: Atelectasis, age: 55, gender: male):

~~~
PREFIX b2v: <http://bio2vec.net/patient_embeddings/function#>
PREFIX BVP: <http://bio2vec.net/patients/BVP_>
PREFIX IMG: <http://bio2vec.net/patients/IMG_>
PREFIX BV: <http://bio2vec.net/patients/>
SELECT ?sim (b2v:similarity(?sim, IMG:00009890_001.png) as ?val) ?p ?f
{
?sim b2v:mostSimilar(IMG:00009890_001.png 10) .
?sim BV:patient ?p .
?sim BV:finding ?f .
}
~~~

The query selects 10 most similar images, of which two of the five most similar ones possess the medical diagnosis *Atelectasis* (patient ID: 8791, Age: 68, Gender: male, similarity: 0.870 and patient ID: 9488, Age: 58, Gender: male, similarity: 0.859). These results are somewhat surprising as the vector extraction employed by the BVLC GoogLeNet is not aimed for any kind of medical image retrieval or analysis, yet nevertheless appears to yield results that indicate the biological relationship.

Furthermore, these results can be enhanced by further constraining the similarity based image retrieval by additional, structured information such as the selection of gender or age ranges within SPARQL. Such query capabilities should allow medical consultants an easier access to relevant cases and facilitate improved diagnosis. It can also enable a faceted exploration of these images and their similarity, using RDF properties as facets to filter on the similarity space. Overall, we also envision our system to be used for other un-structured and semi-structured data in the medical and clinical domain, such as for electrocardiograms (ECGs) and clinical notes.

## 7 Conclusion

Here we presented a first prototype of Vec2SPARQL, a framework which bridges vector space and structured queries in a single endpoint. We have provided a proof of concept for our system using only a single similarity function (cosine similarity) between vectors that can be compared using this function. However, Vec2SPARQL is generic and can be extended with other similarity functions and even functions that have been learned in a supervised manner.

We illustrate the utility of Vec2SPARQL on two use cases: phenotype-driven prioritization of gene–disease associations and retrieval of clinical images based on comparing image feature vectors extracted through deep learning models. Both types of similarity search are combined with structured queries in SPARQL and demonstrate the flexibility and strength of our approach. We believe that Vec2SPARQL serves as a useful framework that can fill the gap between the vector space operations and SPARQL for performing semantic queries on structured and unstructured data.

## Acknowledgements

This work was supported by funding from King Abdullah University of Science and Technology (KAUST) Office of Sponsored Research (OSR) under Award No. URF/1/3454-01-01, FCC/1/1976-08-01, and FCS/1/3657-02-01.

## Footnotes

1 https://github.com/bio-ontology-research-group/vec2sparql

